# Evaluation of variability in high resolution protein structures by global distance scoring

**DOI:** 10.1101/202028

**Authors:** Risa Anzai, Yoshiki Asami, Waka Inoue, Hina Ueno, Koya Yamada, Tetsuji Okada

**Author notes:** Corresponding author: Tetsuji Okada, Department of Life Science, Gakushuin University, 1-5-1 Mejiro, Toshima-ku, Tokyo 171-8588, Japan.

## Abstract

Systematic analysis of statistical and dynamical properties of proteins is critical to understanding cellular events. Extraction of biologically relevant information from a set of high-resolution structures is important because it can provide mechanistic details behind the functional properties of protein families, enabling rational comparison between families. Most of the current structure comparisons are pairwise-based, which hampers the global analysis of increasing contents in the Protein Data Bank. Additionally, pairing of protein structures introduces uncertainty with respect to reproducibility because it frequently accompanies other settings for superimposition. This study introduces intramolecular distance scoring, for the analysis of human proteins, for each of which at least several high-resolution are available. We show that the results are comprehensively used to overview advances at the atomic level exploration of each protein and protein family. This method, and the interpretation based on model calculations, provide new criteria for understanding specific and non-specific structure variation in a protein, enabling global comparison of the dynamics among a vast variety of proteins from different species.

## Introduction

Experimental investigation of protein structures, especially those of human proteins, is an important issue in life science research. The Protein Data Bank (PDB)^1–6^ has recently been growing by approximately 11,000 entries per year^4^; the majority of entries are X-ray structures of human proteins. Conversely, a substantial fraction of PDB entries is archived without accompanying publications. The number of new entries, for each protein with a known structure, is also steadily increasing with efforts applying significant experimental variations. This highlights a need for effective use and comparison of these structures on a global scale.

A detailed and reproducible examination of a set of experimental coordinates is valuable for understanding the structural bases for functionality of a protein, and/or a protein family, at an atomic level. A few structures that differ substantially may be considered to be enough to understand the function of a protein in some cases. However, using most of the available experimental evidence in a global, quantitative, and reproducible manner provides new insights and spurs further advances in structural biology.

Previous work on heptahelical transmembrane (7TM) proteins, such as G protein-coupled receptors (GPCRs), one of the largest protein families in the human genome, suggested that conventional superimposition and root-mean-square deviation (RMSD) calculations are not always optimal for extracting information on the mechanism of transition among the different structural states and on the inherent flexibility^7^. Deviation in a set of structures, possibly originating from specific or non-specific (random) variability of a protein, has not been convincingly quantitated in a manner that makes systematic comparison possible among a large number of proteins. Rmsd is convenient for comparing a limited number of structures, but is pairwise-based, which can result in inconsistent results depending on the algorism of superimposition, and choice of reference. Even the 7TM structures of a GPCR are sometimes superimposed differently, especially with respect to the inactivated and activated forms.

In this work, we aimed to precisely define “crystallographically observed” variation in the intramolecular distances for each protein assigned with a UniProt ID^8^. For example, intramolecular distances between all C_α_ pairs in a polypeptide of 100 amino acids generate 100 × 99 / 2 = 4950 data whereas the number of RMSDs at C_α_s equals to only 100 data. Our previous studies on GPCRs and other 7TM proteins^9^ show that an inverse of “coefficient of variation [standard deviation (stdev) divided by average]”, assigned to the distance of each C_α_ pair, appears as a comprehensive numeral and reasonably reflects the observed variability/invariability; thus, we term it a “score” and term the method as distance scoring analysis (DSA). For example, a score of 100 is equal to 0.1 Å stdev for the distance of 10 Å (with the coefficient of variation of 1%); a higher score indicates higher invariability. Additionally, when all the C_α_ pair scores are averaged for a protein, the value can provide a measure that represents its variability/invariability as observed by X-ray crystallography.

Experimental conditional variations, resulting in multiple PDB entries for a protein, include many factors such as quality of crystals, lattice packing, binding of other proteins and/or small molecules, and engineered mutations. All these factors can induce more or less structural variations, or stabilization in some cases, in a given protein; this can be functionally/inherently specific or non-specific. Thus, when analyzed precisely, a set of X-ray structures, with identically selected initial and terminal positions in a protein, represents the status of the current knowledge on the variability of that protein.

## Results and discussion

The regularly updated statistics in the PDB website indicates that more than a quarter of all entries contain human proteins. Such a prevailing data availability reflects the extensive focus in the structural genomics and makes them a prior subject of our analysis. This work summarizes the results for 300 human proteins, based on the usage of 10,338 PDB entries (the sum of the “entries” column in Supporting Table 1). The statistics for 30 proteins highlighted in the following text are shown in Table 1 The usage ratio to the total entries, archived for these proteins, is 86.3% (91.2% for the 30 proteins in Table 1) [the average of the entries(%) column in the Tables]; this indicates that we have inspected 11,979 entries, which is equal to 36.7% of 32,656 (total X-ray entries for all human proteins as of Aug.28, 2017). The remaining 13.7% (inspected but unused) includes entries containing the protein in the form of such as just a short peptide fragments, or long enough but with lacking (unmodeled) residues. As an initial survey of this long-term project, only continuously modeled X-ray structures of more than 70 residues have been analyzed. Because of stdev calculation, at least three structures are required for a protein to be investigated by DSA. Therefore, among 32,656 entries, roughly more than 5,000 entries of only 1 or 2 entries per protein were not the subject of this study. Also, because of a limitation in the sequence coverage of the archived structures in PDB, the proportion of analyzed length to the full length of a protein varies from 100 to 4.4% [the length(%) columns in the Tables], and the average for 300 proteins is 73.5% (78.5% for the 30 proteins in Table 1). However, even in the partially analyzed proteins, the region covers at least 1 structural domain annotated in Pfam^10^ (**Supporting Table 1**). The most abundant Pfam clan in 300 proteins is Pkinase (18 proteins), followed by NADP_Rossmann (15 proteins) and P-loop_NTPase (12 proteins) clans.

**Table 1.**
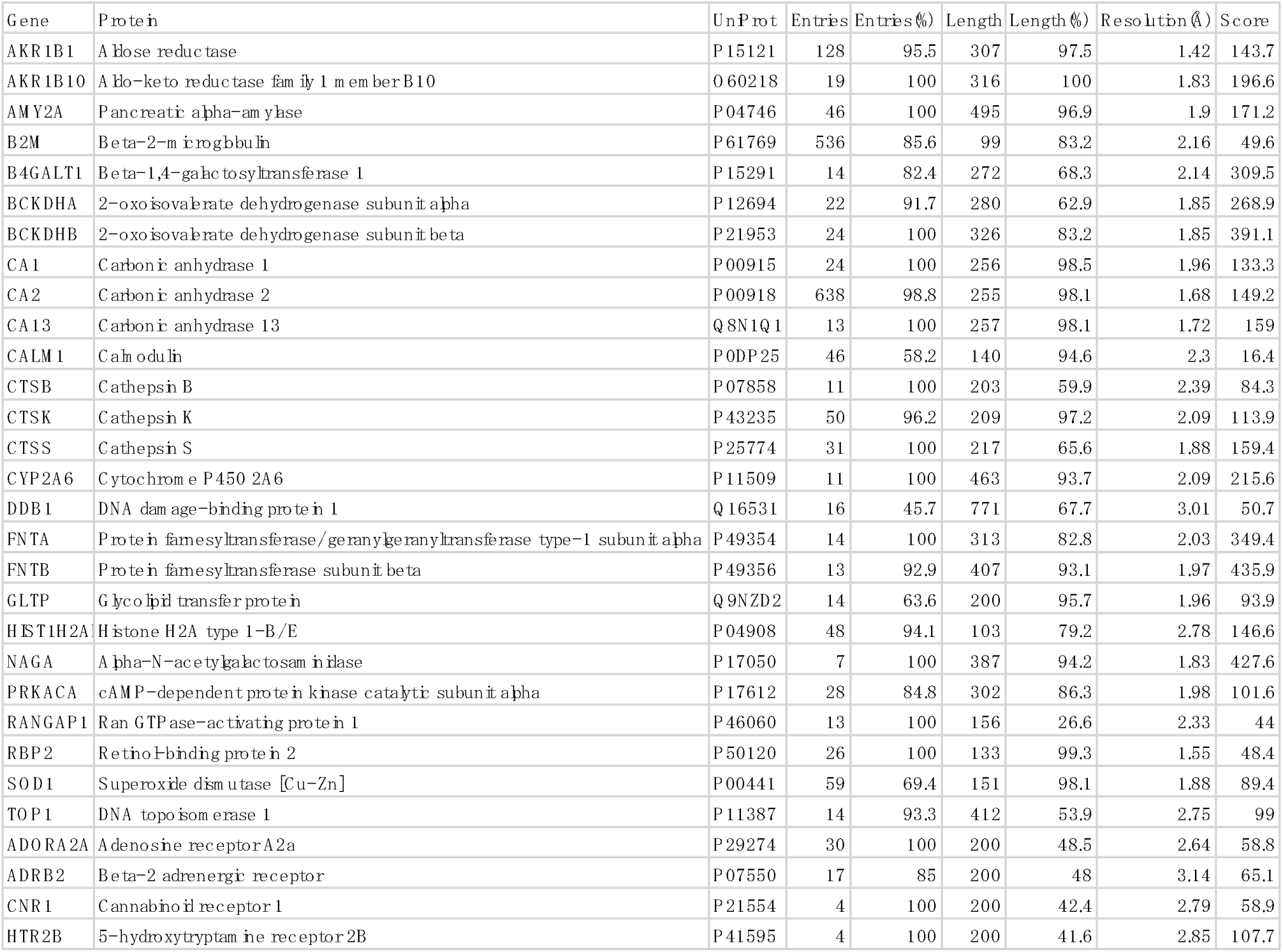
The summary of 30 proteins described in the main text among 300 human proteins analyzed by DSA. Each full-length value, used for calculation of length (%), contains both the signal and activation peptide parts, if they are present.

### The main plot – 2D representation of variability

In Fig. 1, we plotted the score against the average distance for all C_α_ pairs in a protein. Each of the panels is a “main plot” for a protein and the pattern represents the current status of a protein’s 3D structural variation depicted in a 2D figure; this reflects the overall fold and its variability/invariability shown in the lower/upper part of the plot, respectively. The initial purpose of this plot was to delineate the common features in rhodopsin-like GPCRs^7^; then, the usefulness of it was further evaluated with another family of 7TM helical proteins, microbial rhodopsins^9^. Based on these studies, we are confident that the plot can be used for any proteins to concisely grasp crystallographically observed structural variations. In the following, a protein is noted with its primary gene name found in UniProt and the protein name in parenthesis.

**Fig. 1.**
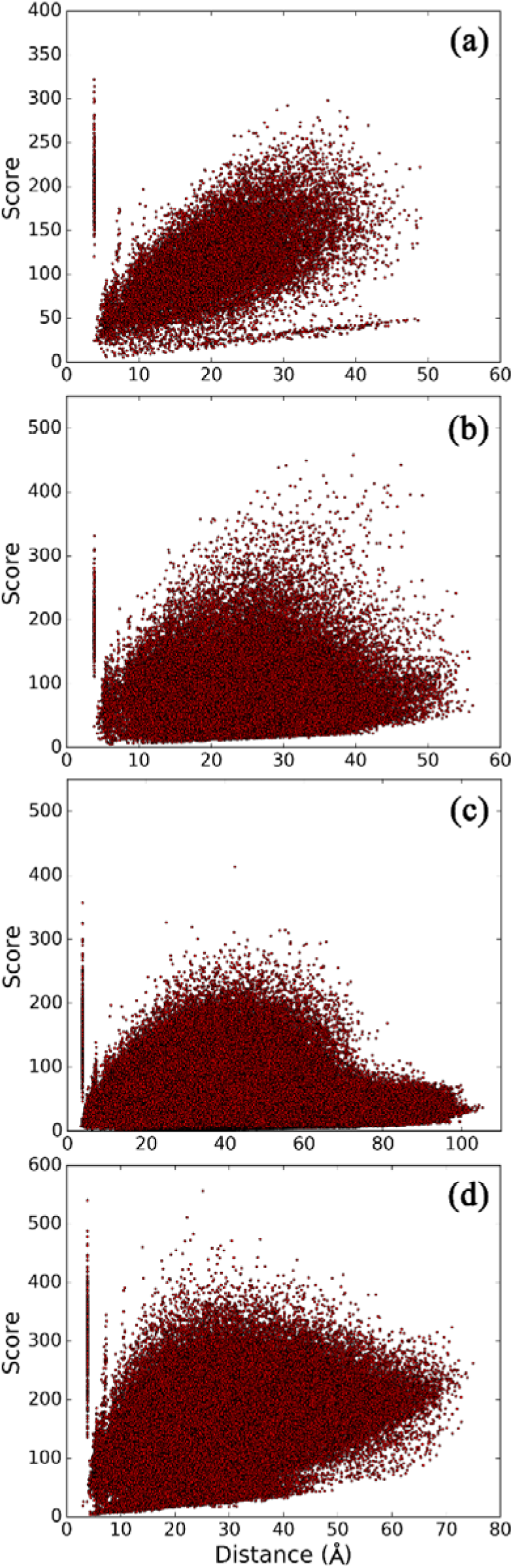
The main plot obtained from distance scoring analysis (DSA). (a) ctsk; (b) prkaca; (c) ddb1; and (d) amy2a.

First, we describe the interpretation of the overall pattern of the plot. In some proteins, the main cluster of the score dots exhibits a round shape lifted toward a longer distance (**Fig. 1a, Supporting Fig. 2a**). The distance dependence is indicative of random and uniform variations in each position of C_α_, because the score increases when stdev is nearly constant at the longer distance. Additionally, some of the proteins exhibiting such a pattern include a thin nearly horizontal array of lower score data points separated from the main body of the plots (**Fig. 1a**). This kind of an array originates from structural variations in some limited part of a polypeptide, such as that in part of the loop region that follows Cys200 linked by a disulfide bond with Cys170 of ctsk (cathepsin K)^11,12^. For all these proteins, the plots indicate that X-ray crystallography has not explored their substantial large-scale structural variations so far.

Among proteins with such a feature, the plot for hist1h2ab (histone H2A.2) is remarkable in terms of that the main body of the data points exhibits a thinner cluster with clearer distance dependency (**Fig. 2c**). Therefore, we attempted to mimic this pattern using model data (**Fig. 2a,b**). The model data was prepared using randomly generated stdev values ranging from 0.05 to 0.4 Å, which were assigned to each of 5253 C_α_ pairs of the 103 residues in the hist1h2ab structures (sequence range 17 to 119). Then, 5253 model scores were calculated using the actual average distance data. To compare the effect of the number of sets of such model scores, we show the two model plots obtained by averaging 3 or 10 sets of artificial 5253 scores. The stdev range of 0.05 to 0.4 Å was chosen based on the minimum and maximum values of the actual stdev data for hist1h2ab.

**Fig. 2.**
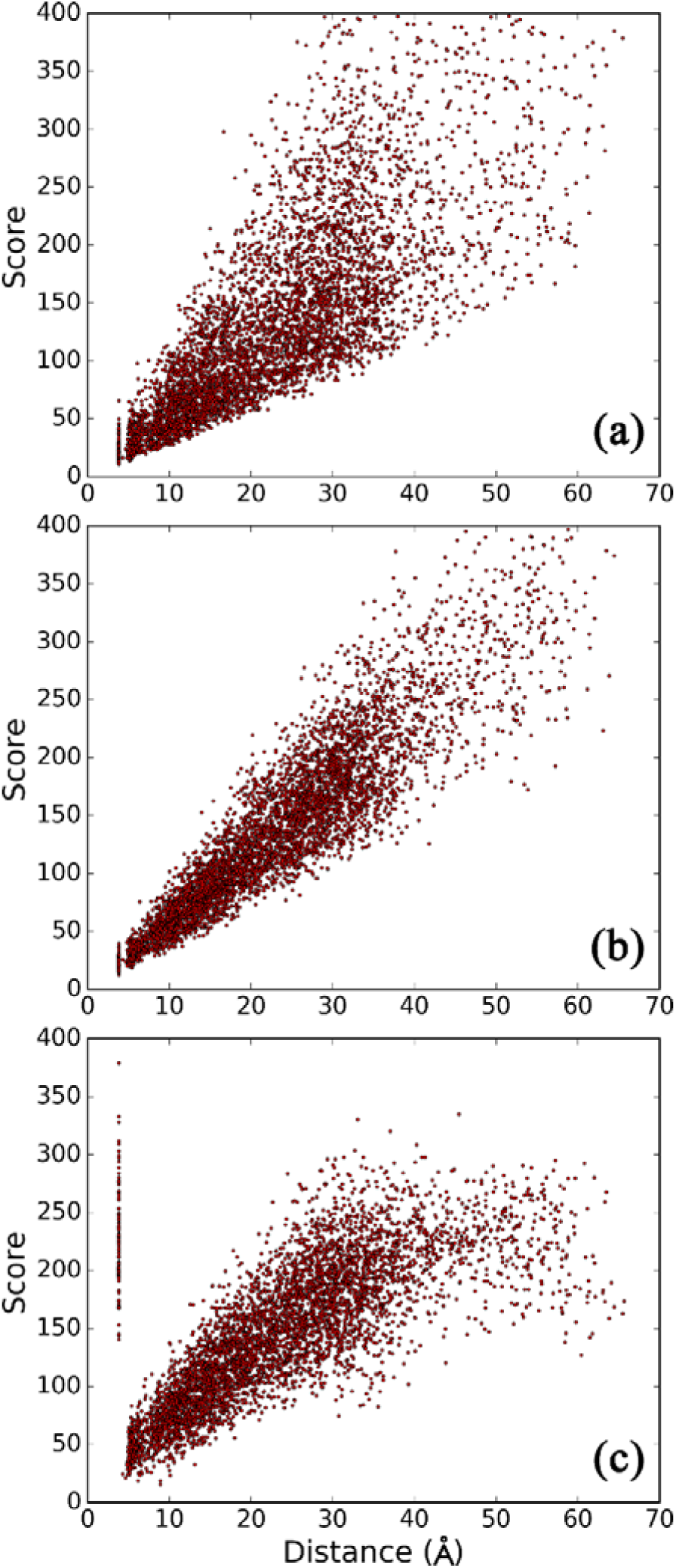
Comparison between model and real main plots of hist1h2ab. (a) Model plot with three random stdev sets; (b) model plot with 10 random stdev sets; (c) real main plot.

From these plots, we suppose that the variation in the analyzed 48 structures of hist1h2ab (**Table 1**) largely arises from random and limited range of deviations in the major part of the examined polypeptide backbone; the large difference between the actual and model data at the longer distance region indicates greater variability in the two terminal parts of the elongated shape of this protein. The main plot pattern for hist1h2ab was found to be an extreme case among 300 proteins. In many cases, the plot exhibits significantly dispersed distribution in the vertical (score) range and without obvious distance dependency; this is shown for prkaca (cAMP-dependent protein kinase catalytic subunit alpha, **Fig. 1b**). It is possible that the unique hist1h2ab pattern represents the fact that the 48 structures were all obtained as a part of a nucleosome^13^, in which hist1h2ab is able to exhibit only a limited range of structural variations.

Interestingly, hist1h2ab, ctsk, and proteins exhibiting a similar profile of the main plot can be grouped as having a common feature; the distribution of the stdev values for all C_α_ pairs in a protein, except for that adjacent to each other (at ~3.8 Å distance), could be well fitted with a log-normal function whereas that of prkaca and many other proteins described below could not (**Supporting Fig. 1**). This observation suggests that such comparison with a model function will provide another distinctive clue on differentiating specific and non-specific structural variations. The log-normal pattern might be explained as that the distance deviations would be somewhat amplified, even if the degree of deviation at each position is similar to each other, as the considered C_α_ pairs in a connected polypeptide chain became apart successively.

In the case of proteins exhibiting large domain movement, a more characteristic main plot pattern appears as represented by ddb1 (DNA damage-binding protein 1, **Fig. 1c**) which is known to rearrange the positioning of its ~300-residue beta-propeller domain against the other part of the polypeptide^14^. There is a clear low-score bump in the main plot of this protein reflecting such a large domain movement. We also found many proteins that exhibit a pattern that is intermediate to those of ctsk and ddb1; this demonstrates that the intramolecular positional variation in a region of a polypeptide manifests as a larger sub-cluster in the main plot when the size of the part increases. In **Supporting Fig.2** we show such pattern changes in the main plot by comparing ctsk with the homologs ctss (cathepsin S) and ctsb (cathepsin B). In this comparison, out of over 200 residues, ctss exhibited scant deviations among 31 structures, whereas ctsb deviated significantly. This is represented in the main plot of ctsb by a low-score sub-cluster which is thicker than ctsk, even in 11 structures, mainly reflecting the high degree of variability in the ~20 residue occluding loop^15,16^.

The plots of other proteins like amy2a (pancreatic α-amylase, **Fig. 1d**) are indicative of outstanding conservation especially at the longest distance region. Expectedly, amy1a (α-amylase 1) and amy2a exhibit a similar overall pattern, despite having a different number of analyzed chains (11 and 48, respectively, **Supporting Fig. 3**). These results show that the main plot reflects global features of a protein, delineated by a set of structures obtained under various conditions.

### The summary plot – global comparison of variability

The “summary plot” (**Fig. 3**) represents a comparison among proteins based on an average score calculated for each protein. For example, the score for akr1b1 (aldose reductase) was calculated as 143.7 by averaging all 46971 C_α_ pair scores obtained from 128 PDB entries, covering 97.5% of the 316 full-length residues (**Table 1**). Because the average X-ray resolution of 128 entries is 1.42, the data point for akr1b1 is shown in the summary plot as x=1.42 and y=143.7 (**Fig. 3c**). The average resolution was set for the x-axis because it was essential to observe whether any substantial score tendency, against quality of the structure data, would appear.

**Fig. 3.**
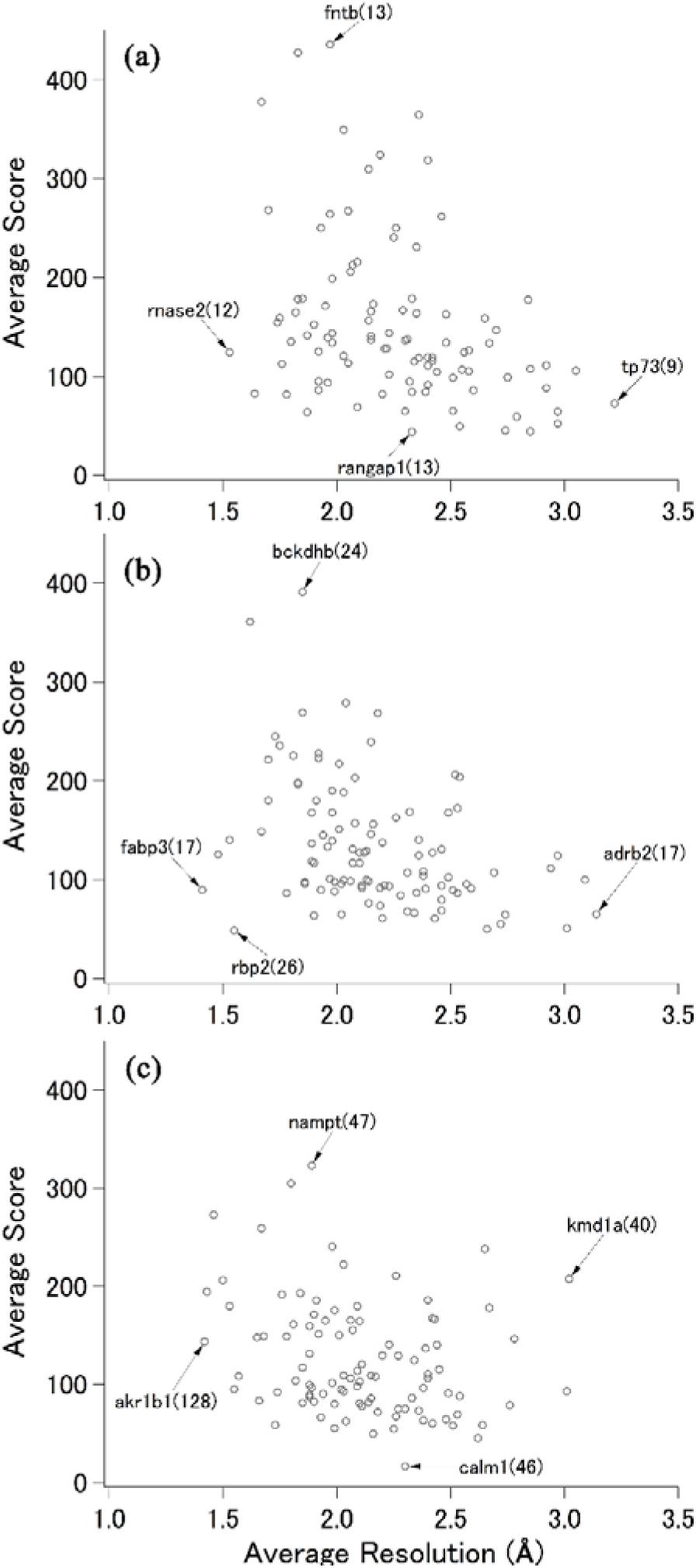
The summary plot obtained from DSA of 300 human proteins. (a) Proteins analyzed using 4~14 structures; (b) proteins analyzed using 15~26 structures; (c) proteins analyzed using 27~638 structures. For each panel, proteins with lowest/highest resolutions and scores are marked by arrows, with the gene name followed by the number of used structures in parentheses.

In Fig. 3, we show scores obtained in this manner for 300 human proteins. The scores are separated into three panels; the top, middle and bottom panels include proteins analyzed using 4 to 14 entries, 15 to 25 entries, and 26 to 638 entries, respectively, including roughly ~100 proteins for each group.

For all panels, there might be only a moderate score dependency against the average resolution (correlation coefficients of −0.35, −0.37, −0.26 for panel a, b, c, respectively). The modest dependency is not surprising because appearance of better quality crystals can become more frequent as the variability of a protein decreases. With respect to dependency on the number of structures used for a protein, the data in the top panel are expectedly enriched for higher score proteins.

Conversely, the scores obtained in all panels are more than 40, with the exception of calm1 (calmodulin, score 16.4). The fact, that 299 out of 300 proteins exhibited average scores of more than 40, indicates that crystallographically observable distance variations, when averaged for all the analyzed C_α_ pairs in a protein, is less than 0.25 Å stdev per 10 Å (2.5%). In other words, proteins that were reluctant to crystallize might exceed this limit because of the presence of highly flexible parts. The score of ~40, in the case of rbp2 (retinol-binding protein 2), is a level that the structure set contains the so-called “domain-swapped” form in a homodimer (PDB ID: 4ZCB). The extensively studied b2m (beta-2-microglobulin), exhibiting the score of less than 50, also shows a domain-swapped form^17^ in the analyzed 536 structures.

The average score for each protein correlated well with the type of Pfam clan to which the analyzed domain belongs. The scores of 10 out of 12 proteins in the P-loop_NTPase clan were less than 87, whereas all 12 proteins in the NADP_Rossmann clan exhibit scores of more than 99. More specifically, the scores of three carbonic anhydrases are in the narrow range of 132 ~ 160, and we confirmed this kind of score similarity in many types of homologous protein groups, as shown in Supporting Table 1. The average score for 300 proteins was ~137. The highest scoring protein was fntb (protein farnesyltransferase subunit beta) with the score of 435.9; the cognate heterodimerization partner fnta also exhibited a high score (349.4). The second and third were naga (alpha-N-acetylgalactosaminidase) and bckdhb (mitochondrial 2-oxoisovalerate dehydrogenase subunit beta) with the scores of 427.6 and 391.1, respectively. The structures of bckdhb were also solved in a heterodimer with bckdha, which exhibited the score of 268.9. A survey of such high-scoring proteins suggests that two factors, one biological and the other technical, may be involved in retaining the high score: tight association between long polypeptides and structure determination in a single crystal lattice type, respectively. With regard to the latter, any proteins solved in a single lattice exhibit low scores, provided that any conditional variations other than lattice packing successfully provoked substantial structural deviations. On the other hand, the scores shown in the present study for some of the high-scoring proteins can decrease substantially as the availability of crystal structures solved in multiple lattice types for each of them increases. Importantly, the present scoring helps recognizing upper and lower limits of the structural variability of the functionally folded polypeptides.

For exploring another global tendency of score variability, the x-axis of the summary plot can represent any parameters other than the average resolution; such parameters can be the length of polypeptide, percentage of alpha-, and beta-folds. Domain scores can also be examined including many “partial entries” that have not been used in the present study.

### The progress plot

Another data tool, useful for interpreting the average score of a protein, is the “progress plot” (**Fig. 4**). This plot is used to examine how the average score (the y-value of each point in Fig. 3) converges and/or changes upon the addition of structure (PDB entry). In this work, only one chain per entry was used for a protein in order to avoid possible bias from including, for example, symmetry-averaged very similar structures. Thus, the number of chains (indicated by the x-axis in Fig. 4) is identical to the number of X-ray entries, for a continuously modeled protein, used for analysis with defined initial and final residues. The order of chain addition (from the left to the right, along the x-axis in Fig. 4) corresponds to the historical progress, in most cases, because we analyzed data in a filename-sorted manner, which appears to match the recent PDB style of archiving. Thus, the initial score, at the number of chains = 3, indicates the degree of variation among the first three name-sorted structures for a protein; the values obtained for 300 proteins vary from less than 50 for calm1 to over 1000 for nampt (nicotinamide phosphoribosyltransferase).

**Fig. 4.**
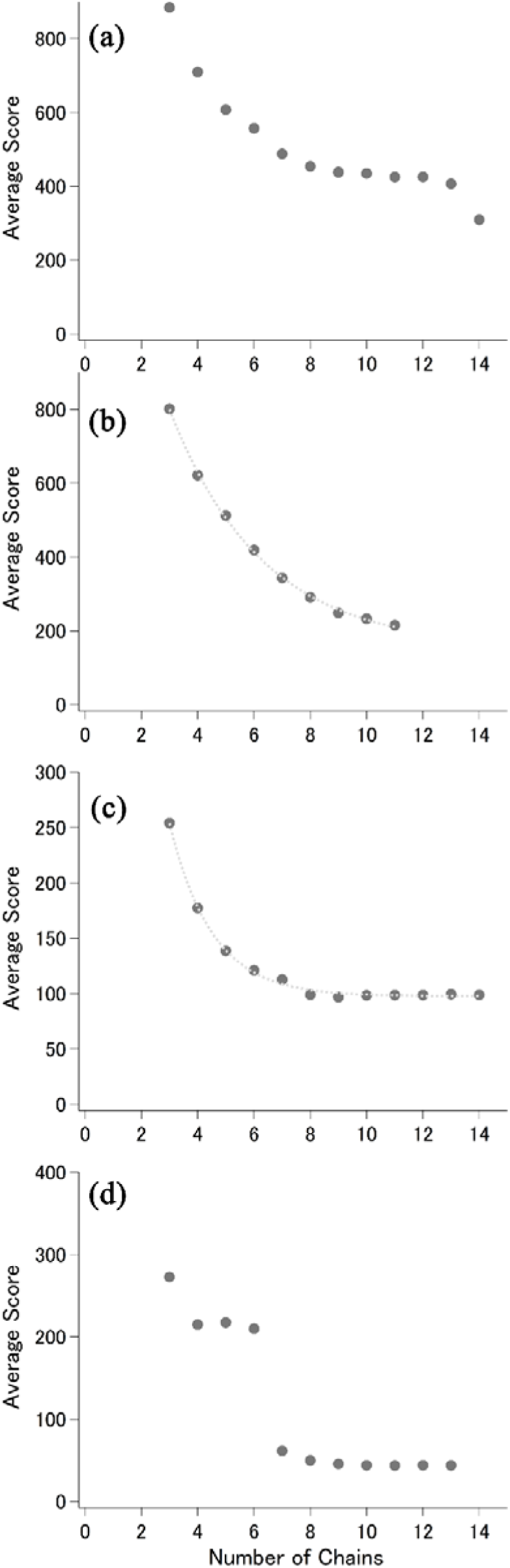
Progress plot obtained from DSA. (a) b4galt1; (b) cyp2a6; (c) top1; and (d) rangap1. For cyp2a6 and top1, exponential fittings are overlaid with dotted curves.

Here we show four examples of the progress plot; two of these (cyp2a6 and top1) represent seemingly non-specific variations, and the other two (b4galt1 and rangap1) show distinct structural states. In Fig. 4 a~d, these proteins are arranged in the order of higher score to lower score [309.5, 215.6, 99.0, and 44.0 for b4galt1, cyp2a6, top1, and rangap1, respectively, at the final (right end) value].

These plots will help elucidate how the average score of a protein converges and/or varies. The data on cyp2a6 (cytochrome P450 2A6) and top1 (DNA topoisomerase 1) were well-fitted with single exponentials. Such exponential convergence, upon the addition of structures, can be simulated using random distances, varying from the observed average to the maximum and the minimum, among the experimental values for each C_α_ pair. When we performed this calculation for hist1h2ab, using the average/maximum/minimum data from 48 structures, the average score nearly converged with less than 10 structures, whereas the actual progress plot for this protein exhibited more irregular and moderate convergence (**Supporting Fig. 4a**). Therefore, random sampling of the possible largest distance range can result in the convergence of the average score with several structures in a single-exponential approach.

The significant and discrete score changes (**Fig. 4a, d**), found in b4galt1 (beta-1,4-galactosyltransferase 1) at the number of chains from 13 to 14, and rangap1 (Ran GTPase-activating protein 1) at the number of chains from 5 to 6, can be explained by the inclusion of structures solved in complex with the respective binding partners (lactoalbumin for b4galt1 and SUMO-1 for rangap1^18^); this correspondence was made by referencing the “entry table”, which was prepared for each protein and summarized the details of all entries (title, space group, lattice constants, etc.) used for DSA. An example of the entry table for b4galt1 is shown in Supporting Table 2, and the remaining tables for 299 proteins will be made available from our web site (gses.jp)

Additional interesting results were obtained from a comparison of three carbonic anhydrases analyzed in this study (**Table 1**). Ca1 and ca13 exhibited monotonous convergence that was similar to each other, resulting in the current score of 133.3 with 24 structures and 159.0 with 13 structures, respectively (**Supporting Fig. 4b, d**). In contrast, ca2 initially converged to the score of over 200; this score reached the level of ca1 and ca13 only after exceeding ~65 entries (**Supporting Fig. 4c**). Again, from the entry table, it is possible that the difference between ca1/ca13 and ca2 may arise from how lattice variations are involved in each of the structure sets. The first ~60 entries of ca2 data are solved in an identical space group with similar lattice constants^19^, whereas the data sets for ca1 and ca13 are each composed of more than three types of lattices. This example shows not only that homologous proteins converge to reasonably similar scores but that convergence can be reached experimentally with several structures. We found many other proteins, for which nearly full structural divergence has been explored, represented by low scores with an even fewer number of chains. A recent and clear example of such a case is cannabinoid receptor 1 (cnr1), a rhodopsin-like GPCR, which exhibited a score of 58.9, calculated with a data set containing only four chains and reflecting the fact that the set includes both inactivated and activated forms^20,21,22^.

### All-alpha proteins

To gain further insights from the main plot of each protein, and from comparison among proteins, we examined all-alpha proteins and the short distance range (**Fig. 5**). In addition to 27 proteins annotated in PDB as having all-alpha folds of the SCOP category^23^, 5 GPCRs were included in 300 proteins shown in Fig. 3. In the distance range from 3 to 8 Å, two features were obvious.

**Fig. 5.**
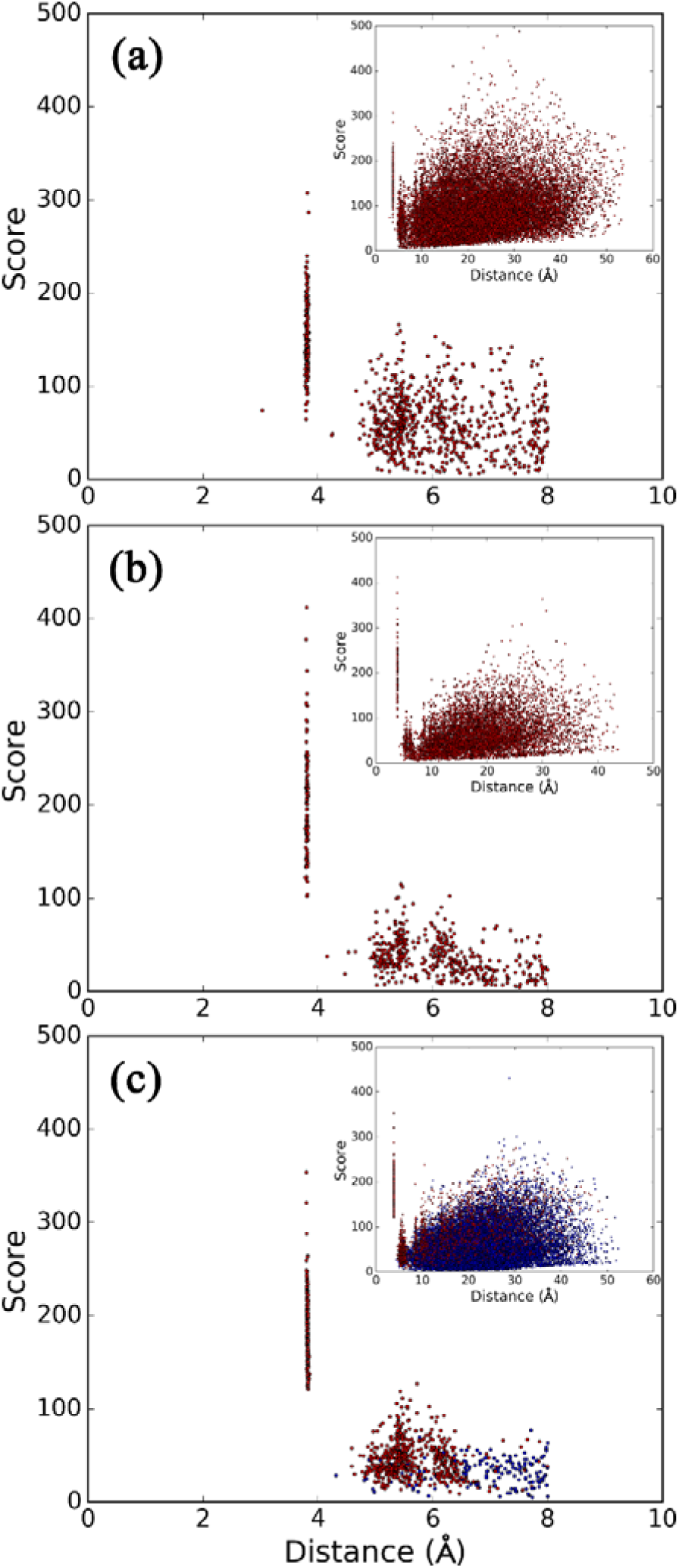
The main plot of all-alpha proteins focusing on short-distance range. (a) gltp; (b) il4; (c) adrb2. Inset: whole view of the main plot For adrb2, intrahelical and interhelical C_α_ pairs are colored in red and blue dots, respectively.

The first feature, which is common to all proteins, regardless of the type of fold, is a nearly straight vertical array of data points at ~3.8 Å, originating mostly from adjacent C_α_ pairs. In addition, many proteins exhibit a few points at distances shorter than this array, indicating the presence of *cis* peptide linkages. As shown in Fig. 1d (amy2a) and Fig. 5a (gltp, glycolipid transfer protein), the scores of such points are frequently lower than those of most C_α_ pairs constituting the ~3.8 Å array; this suggests substantial inconsistency with regard to the *cis/trans* peptide linkage, mostly involving a proline, for the pair position among the structures. Because the assignment of these *cis*-containing adjacent pairs, in the context of 3D structure, is easily accomplished, further detailed global analysis will be useful. The second feature is a thicker cluster of data points in the distance range from ~5 to ~7 Å, which is clearer in all-alpha proteins; it can be further separated into two clusters as in the case of il4 (interleukin 4, **Fig. 5b**). When we look at these data, and those of GPCRs (**Fig. 5c**), analyzed only for the 7TM helices (excluding all interhelical loops and two terminal tails), a vacant region between the two features becomes obvious. This observation indicates that any of the interhelical C_α_-C_α_ distances rarely exhibit distances at around 4.5 Å. Conversely, we found a consistent interhelical C_α_ pair in 3 of the 4 rhodopsin-like GPCRs (adora2a, adrb2, htr2b) exhibiting the average distance of less than 4.5 Å. Because we have been studying the 7TM helical core region of GPCRs, assignment of data points in the main plot to the structural part was also easily accomplished. The consistent pair consisted of one amino acid from helix I (1.46 by BW numbering^24^) and the other from helix VII (7.47), indicating a common backbone contact in these receptors. The residue at 7.47 is located close to the NPxxY motif, which is important in the activation process of rhodopsin-like GPCRs^25–27^. These results demonstrate how we can extract detailed information for a protein, or a set of proteins from the main plot of DSA.

### Implications from high scoring pairs

Systematic appearance of high scores, in the upper region of the main plot, can be an indication of a specifically restrained region and/or direction within a protein. Although the score for a single C_α_ pair, obtained from a set of several structures is not enough for a sufficient level of confidence, we frequently observed that a few specific residues and/or regions contribute to the majority of the highest scores in a protein. Fig. 6 shows the examples of such observations. In akr1b1, which has been analyzed using more than 100 structures, the highest scoring pairs, in the distance range from 10 to 25 Å are represented by the residues in parts of beta strands occupying the core of the protein (**Fig. 6a**). Because this analysis covers 97.5% of the full-length akr1b1, it is likely that this result represents the inherent invariability of this region. Similar results were obtained for akr1b10 using only 19 structures. In the other case, as shown for sod1 (superoxide dismutase 1) analyzed using 59 structures, a peripheral part of the 3D structure was highlighted as exhibiting high scores, mainly involving the residues in the C-terminal strand (**Fig. 6b**). Interestingly, this strand was recently implicated as one of the aggregation-triggering segments^28,29^. The result on lyz (lysozyme, **Fig. 6c**) also highlighted a beta-strand, which includes an amyloidogenic mutation site (Ile56) ^30,31^, as exhibiting high scores.

**Fig. 6.**
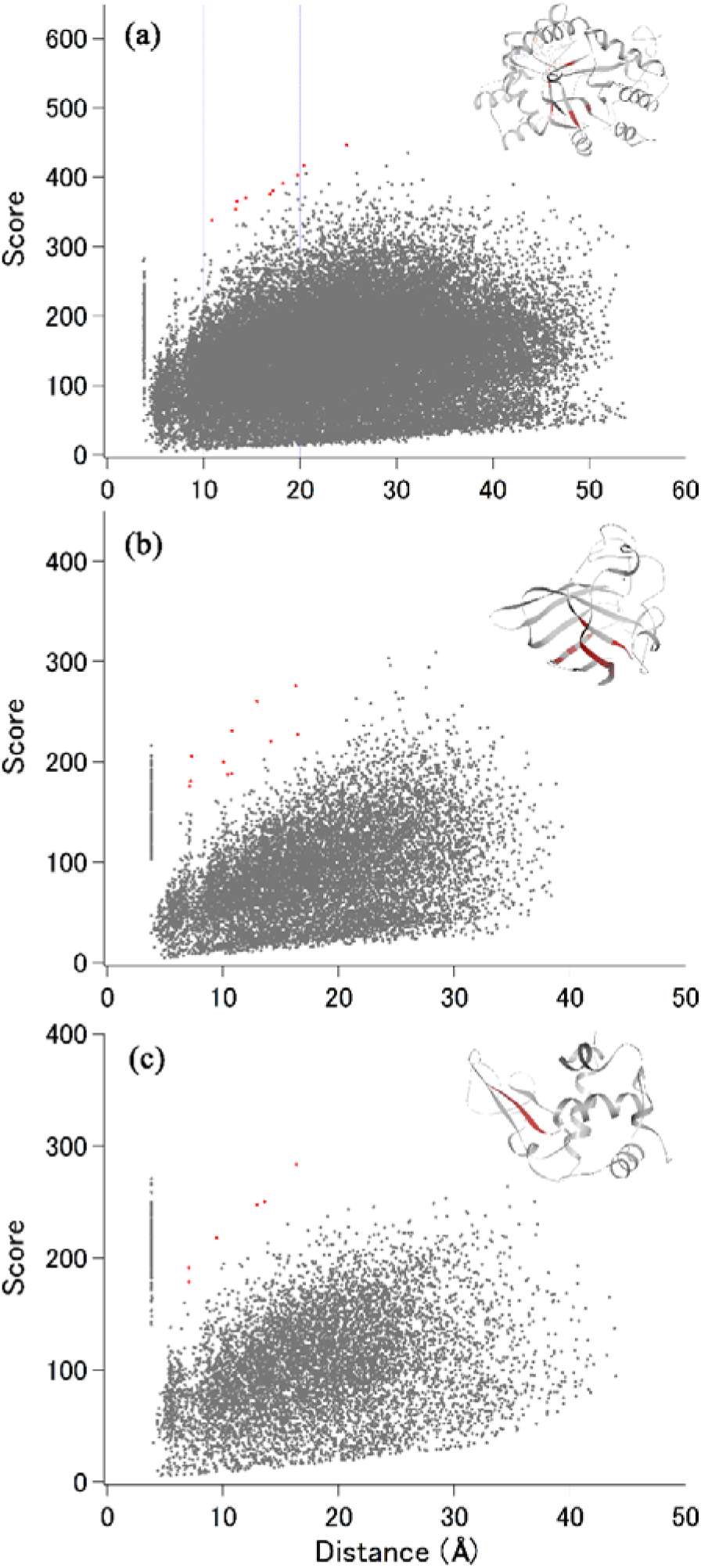
The main plot highlighting systematic appearance of high-scoring pairs. (a) akr1b1; (b) sod1; (c) lyz. All the residues contributing to the red dots in the main plot are graphically shown in the inset as red part of the ribbon. The PDB ID of the inset is (a) 4YU1; (b) 5U9M (chain A); (c) 208L.

The amount of high scoring pairs, clearly distinct from the main cluster, can be scarce depending not only on the type of protein but also on the distance range in a protein (**Fig. 1**). Conversely, a substantial amount of beta folds in a protein tends to exhibit an array of data points in the main plot, frequently reaching high scores at the distance of ~7 Å (**Fig. 1a, d**), which corresponds to the distance between residue numbers i and i+2 in a strand. Further detailed analysis of high scoring pairs, in short or long distance ranges, will be beneficial.

In summary, DSA, and the derived plots and tables described above, will help summarize the progress and current status of X-ray studies for a number of proteins and cross comparison among proteins. This approach will also provide valuable information, not readily discernible by other methods, on each protein and protein family. On the other hand, some proteins described in this study, such as fnta and prkaca, have mammalian orthologs that have been more extensively studied by X-ray crystallography. Our approach can be extended to the structural compassion of structures from different species as well as a comparison of structures between the species. Thus, further extension, improvement, and updating of this method will contribute to the community of structural and molecular biology.

## Methods

### Data selection

After preliminary analyses of proteins, such as the T4 lysozyme and hras (GTPase HRas), optimizing data manipulation procedure, human protein PDB entries were searched and downloaded from August, 2016 to August, 2017. To survey all the human X-ray entries, we constructed a table based on Custom Reports from PDB, which resulted in a list of over 70,000 structures. This table was then sorted by UniProt ID^8,32^ and used to prepare an additional table containing information on the group of candidates for DSA, such as the name of the protein, number of amino acids, and X-ray entries per unique UniProt ID of the protein; this table was then sorted by the amount of X-ray entries. Thereby, we obtained 1,415 proteins with more than four entries each. For the 300 proteins analyzed in this study, the order of entries, and the range of the primary sequence of the chain, processed by the score-analyzer (see below), were also tabulated (“entry table” described in the main text). The selection of 300 proteins (**Fig. 3, Supporting Table 1**) was random; however, we attempted to cover a wide range of the chain length and resolution.

The 7TM GPCR structures for DSA were obtained from our archive at gses.jp/7tmsp. Each of the chains contains 200 amino acids, defined previously^33^ for reproducible comparison of all receptors including non-rhodopsin-like GPCRs.

### Distance scoring analysis procedure

A modified version of a Python script (score-analyzer^9^, sa_v16) performs the following functions: reading a PDB entry, extracting/displaying of the C_α_ coordinates (choosing one from the alternative conformations, if present), calculating and storing distance data per chain for a selected sequence range, calculating average scores for making a “progress plot”, extracting and assessing the high-scoring populations in a defined set of distance ranges, and plotting and saving graphs and tables. Additional functions for GPCRs are implemented to associate the general amino acid position numbers (BW numbers)^24^ with each C_α_ and to classify the C_α_–C_α_ pairs (e.g., intrahelical or interhelical, **Fig. 5c**). For a protein having a unique UniProt ID and more than four X-ray PDB entries (more than three in the case of GPCRs), a sequence range of C_α_ coordinates was determined by referencing the “Protein Feature View of PDB entries mapped to a UniProtKB^8^ sequence” page in the PDB; all the C_α_–C_α_ distances in this range were calculated per chain. For GPCRs, 200-residue 7TM coordinates, archived on our web site (gses.jp/7tmsp), were used without any selection of sequence range. Due to the lack of polypeptides (because of unmodeled amino acids), not all entries were investigated; percentages of processed entries are shown in Table 1 and Supporting Table 1. All the extracted C_α_ in a chain were inspected using the main window of sa_v16, ensuring that neither break nor duplication was present, and that one C_α_ was extracted per amino acid position in a sequence. The distance data were stored, and the procedure was repeated for the entries in a selected set, ensuring again that no inconsistency in the positioning of C_α_–C_α_ pairs was present in any of the chains, using the main-table window of sa_v16. The “progress plot” against the number of chains (**Fig. 1C**) was obtained at this stage. Then, the final scores for each C_α_–C_α_ pair calculated using all the available entries were plotted against the respective average distances (“main plot”), and the specific distance and/or score ranges were examined further (**Figs. 5, 6**). Fitting of the stdev distribution with Gaussian and log-normal curves (**Supporting Fig. 1**) and of the progress plot with exponentials (**Fig. 4 and Supporting Fig. 4**) was conducted using Igor Pro (WaveMetrix).

Most of the plots were prepared with matplotlib, implemented in sa_v16; other plots were drawn using Igor Pro. Protein graphics were prepared using the DS Visualizer (Biovia).

## Acknowledgments

Supported by the Ministry of Education, Culture, Sports, Science and Technology of Japan.

## Contributions

R.A, Y.A., W.I., H.U., K.Y., and T.O. performed the analysis, T.O. designed the study and wrote the manuscript.

